# Rapid Core-Genome Alignment and Visualization for Thousands of Intraspecific Microbial Genomes

**DOI:** 10.1101/007351

**Authors:** Todd J. Treangen, Brian D. Ondov, Sergey Koren, Adam M. Phillippy

## Abstract

Though many microbial species or clades now have hundreds of sequenced genomes, existing whole-genome alignment methods do not efficiently handle comparisons on this scale. Here we present the Harvest suite of core-genome alignment and visualization tools for quickly analyzing thousands of intraspecific microbial strains. Harvest includes Parsnp, a fast core-genome multi-aligner, and Gingr, a dynamic visual platform. Combined they provide interactive core-genome alignments, variant calls, recombination detection, and phylogenetic trees. Using simulated and real data we demonstrate that our approach exhibits unrivaled speed while maintaining the accuracy of existing methods. The Harvest suite is open-source and freely available from: http://github.com/marbl/harvest.

## Rationale

Microbial genomes represent over 93% of past sequencing projects, with the current total over ten thousand and growing exponentially. Multiple clades of draft and complete genomes comprising hundreds of closely related strains are now available from public databases [1], largely due to an increase in sequencing-based outbreak studies [2]. The quality of future genomes is also set to improve as short-read assemblers mature [3] and long-read sequencing enables finishing at greatly reduced costs [4,5].

One direct benefit of high-quality genomes is that they empower comparative genomic studies based on multiple genome alignment. Multiple genome alignment is a fundamental tool in genomics essential for tracking genome evolution [6–8], accurate inference of recombination [9–14], identification of genomic islands [15,16], analysis of mobile genetic elements [17,18], comprehensive classification of homology [19,20], ancestral genome reconstruction [21], and phylogenomic analyses [22–24]. The task of whole-genome alignment is to create a catalog of relationships between the sequences of each genome (ortholog, paralog, xenolog, etc. [25]) to reveal their evolutionary history [26,27]. While several tools exist (LS-BSR [28], Magic[29], Mavid[30], Mauve [31–33], MGA [34], M-GCAT [35], Mugsy [36], TBA [37], multi-LAGAN [38], PECAN [39]), multiple genome alignment remains a challenging task due to the prevalence of horizontal gene transfer [40,41], recombination, homoplasy, gene conversion, mobile genetic elements, pseudogenization, and convoluted orthology relationships [25]. In addition, the computational burden of multiple sequence alignment remains very high [42] despite recent progress [43].

The current influx of microbial sequencing data necessitates methods for large-scale comparative genomics and shifts the focus towards scalability. Current microbial genome alignment methods focus on all-versus-all progressive alignment [31,36] to detect subset relationships *(i.e.* gene gain/loss), but these methods are bounded at various steps by quadratic time complexity. This exponential growth in compute time prohibits comparisons involving thousands of genomes. Chan and Ragan [44] reiterated this point, emphasizing that current phylogenomic methods, such as multiple alignment, will not scale with the increasing number of genomes, and that “alignment-free” or exact alignment methods must be used to analyze such datasets. However, such approaches do not come without compromising phylogenetic resolution [45].

Core-genome alignment is a subset of whole-genome alignment, focused on identifying the set of orthologous sequence conserved in all aligned genomes. In contrast to the exponential complexity of multiple alignment, core-genome alignment is inherently more scalable because it ignores subset relationships. In addition, the core genome contains essential genes that are often vertically inherited and most likely to have the strongest signal-to-noise ratio for inferring phylogeny. The most reliable variants for building such phylogenies are single-nucleotide polymorphisms (SNPs). Thus, core-genome SNP typing is currently the standard method for reconstructing large phylogenies of closely related microbes [46]. Currently, there are three paradigms for core-genome SNP typing based on read mapping, k-mer analyses, and whole-genome alignment.

Read-based methods have dominated the bioinformatics methods landscape since the invention of high-fidelity, short-read sequencing (50-300bp) [47]. This has made it very affordable to sequence, yet extremely challenging to produce finished genomes [48,49]. Thus, comparative genomics has turned to highly efficient and accurate read mapping algorithms to carry out assembly-free analyses, spawning many mapping tools [50-53] and variant callers [54-56] for detecting SNPs and short Indels. Read-based variant calling typically utilizes a finished reference genome and a sensitive read mapper (BWA [52], Smalt), variant caller (samtools/bcftools [56], GATK [54]), and variant filter (minimum mapping quality, core genomic regions). This method has been shown effective in practice [57] and does not rely on assembly. However, mapping requires the read data, which is not always available and can be orders of magnitude larger than the genomes themselves. In addition, mapping can be sensitive to contaminant, overlook structural variation, misalign low-complexity and repetitive sequence, and introduce systematic bias in phylogenetic reconstruction [58–60].

Exact alignment methods, often formulated as k-mer matching, can produce high precision results in a fraction of the time required for gapped alignment methods [63]. Spectral k-mer approaches have been used to estimate genome similarity [64], and k-mer based methods are commonly used to identify or cluster homologous genomic sequence [65,66]. Recently, k-mers have also been extended to SNP identification. kSNP [67] identifies odd-length k-mers between multiple samples that match at all but the central position. The matched k-mers are then mapped back to a reference genome to locate putative SNPs. Conveniently, this approach is suitable for both assembled genomes and read sets, but sensitivity is sacrificed for the improved efficiency of exact alignment [68].

Genome assembly [69–77], followed by whole-genome alignment [38,78, 79], is the original method for variant detection between closely related bacterial genomes [80] and has been shown to perform well across multiple sequencing platforms [81]. In addition to SNPs, whole-genome alignment is able to reliably identify insertions and deletions (Indels) and other forms of structural variation. Thus, whole-genome alignment is the gold standard for comprehensive variant identification, but relies on highly accurate and continuous assemblies, which can be expensive to generate.

Lastly, and unlike reference mapping, whole-genome alignment is not easily parallelized or scaled to many genomes.

Specifically for the task of whole-genome SNP typing, the choice of read- or genome-based methods can often depend on data availability. For example, of the 24,000 bacterial genomes currently in NCBI RefSeq [82], only 55% have associated SRA read data and analysis of the remaining 45% requires genome-based methods. Thankfully, recent advances in both sequencing technology and assembly algorithms are making microbial genomes more complete than ever before. Modern *de Bruijn* assemblers like SPAdes [83]are able to generate high-quality assemblies from short reads [3], and long read technologies have enabled the automated finishing of microbial genomes for under $1,000 [84]. With the number of publically available genomes currently doubling every 18 months [1], and genome quality improving with the arrival of new technologies, we set out to solve the problem of aligning thousands of closely-related whole genomes.

## Rapid Core-Genome Alignment and Visualization

Here we present Parsnp and Gingr for the construction and interactive visualization of massive core-genome alignments. For alignment, Parsnp combines the advantages of both whole-genome alignment and read mapping. Like whole-genome alignment, Parsnp accurately aligns microbial genomes to identify both structural and point variations, but like read mapping, Parsnp scales to thousands of closely related genomes. To achieve this scalability, Parsnp is based on a suffix graph data structure for the rapid identification of maximal unique matches (MUMs), which serve as a common foundation to many pairwise [78,79,85] and multiple genome alignment tools [31–36]. Parsnp uses MUMs to both recruit similar genomes and anchor the multiple alignment. As input, Parsnp takes a directory of MultiFASTA files to be aligned; and as output, Parsnp produces a core-genome alignment, variant calls, and a SNP tree. These outputs can then be visually explored using Gingr. The details of Parsnp and Gingr are described below.

### MUMi Recruitment

Parsnp is designed for intraspecific alignments and requires input genomes to be highly similar (e.g. within the same subspecies group or >=97% average nucleotide identity). For novel genomes or an inaccurate taxonomy, which genomes meet this criterion is not always known. To automatically identify genomes suitable for alignment, Parsnp uses a recruitment strategy based on the MUMi distance [86]. Only genomes within a specified MUMi distance threshold are recruited into the full alignment.

### Compressed Suffix Graph

Parsnp utilizes a Directed Acyclic Graph (DAG) data structure, called a Compressed Suffix Graph (CSG), to index the reference genome for efficient identification of multi-MUMs. CSGs have the unique property of representing an optimally compressed structure, in terms of number of nodes and edges, while maintaining all intrinsic properties of a Suffix Tree. CSGs were originally proposed as a more space-efficient alternative to Suffix Trees and first implemented in M-GCAT [35]. Node and edge compression of the Suffix Tree incurs a linear-time construction penalty, but facilitates faster traversal of the structure once built. Provided sufficient memory, the CSG can be used to align genomes of any size; however, the current implementation has been optimized for microbial genomes, requiring approximately 32 bytes per reference base for CSG construction and 15 bytes per base for the aligned genomes.

Note that because multi-MUMs are necessarily present in all genomes, the choice of a reference genome has no effect on the resulting alignment.

### Multi-MUM Search

Once built for the reference genome, all additional genomes are streamed through the CSG, enabling rapid, linear-time identification of MUMs shared across all genomes. A divide-and-conquer algorithm, adapted from M-GCAT [35], recursively searches for smaller matches and iteratively refines the multi-MUMs. Next, locally collinear blocks (LCBs) of multi-MUMs are identified. These LCBs form the basis of the coregenome alignment.

### Parallelized LCB Alignment

The multi-MUMs within LCBs are used to anchor multiple alignments. Gaps between collinear multi-MUMs are aligned in parallel using MUSCLE [87]. To avoid the unnecessary overhead of reading and writing MultiFASTA alignment files, Parsnp makes direct library calls via a MUSCLE API. The MUSCLE library is packaged with Parsnp, but originally sourced from the Mauve code base [88]. As with Mauve, MUSCLE is used to compute an accurate gapped alignment between the match anchors. Though MUSCLE alignment can be computationally expensive, for highly similar genomes, the gaps between collinear multi-MUMs are typically very short (e.g. a single SNP column in the degenerate case).

### SNP Filtering and Trees

The final Parsnp multiple alignment contains all SNP, Indel, and structural variation within the core genome. However, given their ubiquity in microbial genome analyses, Parsnp performs additional processing of the core-genome SNPs. First, all polymorphic columns in the multiple alignment are flagged to identify: (i) repetitive sequence, (ii) small LCB size, (iii) poor alignment quality, (iv) poor base quality, and (v) possible recombination. Alignment quality is determined by a threshold of the number of SNPs and Indels contained within a given window size. Base quality is optionally determined using FreeBayes [55] to measure read support and mixed alleles. Bases likely to have undergone recent recombination are identified using PhiPack [89]. Only columns passing a set of filters based on these criteria are considered reliable core-genome SNPs. The final set of core-genome SNPs is given to FastTree2 [90] for reconstruction of the whole-genome phylogeny.

### Compressed Alignment File

For simplicity and storage efficiency, the output of Parsnp includes a single binary file encoding the reference genome, annotations, alignment, variants, and tree. Thousandfold compression of the alignment is achieved by storing only the columns that contain variants. The full multiple alignment can be faithfully reconstructed from this reference-compressed representation on demand. Since Parsnp focuses on aligning only core blocks of relatively similar genomes, the number of variant columns tends to increase at a sub-linear rate as the number of genomes increases, resulting in huge space savings versus alternative multiple alignment formats. Conversion utilities are provided for importing/exporting common formats to/from the binary archive file, including: BED, GenBank, FASTA, MAF, Newick, VCF, and XMFA.

### Interactive Visualization

Developed in tandem with Parsnp, the visualization tool Gingr allows for interactive exploration of trees and alignments. In addition to the compressed alignment format, Gingr accepts standard alignment formats and can serve as a general-purpose multiple alignment viewer. Uniquely, Gingr is capable of providing dynamic exploration of alignments comprising thousands of genomes and millions of alignment columns. It is the first tool of its kind capable of dynamically visualizing multiple alignments of this scale. The alignment can be seamlessly zoomed from a display of variant density (at the genome level) to a full representation of the multiple alignment (at the nucleotide level). For exploration of phyletic patterns, the alignment is simultaneously presented along with the core-genome SNP tree, annotations, and dynamic variant highlighting. The tree can be zoomed by clade, or individual genomes selected to expand via a fisheye zoom. Structural variation across the genome can also be displayed using Sybil coloring [91], where a color gradient represents the location and orientation of each LCB with respect to the reference. This is useful for identifying structurally variant regions of the core.

## Evaluation of Performance

We evaluated Parsnp on three simulated datasets (derived from *Escherichia coli* K-12 W3110) and three real datasets *(Streptococcus pneumoniae, Peptoclostridium difficile, Mycobacterium tuberculosis).* Parsnp is compared below versus two whole-genome alignment methods (Mugsy, Mauve), a k-mer based method (kSNP), and two commonly used mapping pipelines (based on Smalt and BWA). The Smalt pipeline replicates the methods of the landmark Harris *et al.* paper [92] that has been adopted in many subsequent studies. The BWA pipeline is similar to the Smalt pipeline, but uses BWA for read mapping (Methods).

### Simulated Escherichia coli W3110 Dataset

To precisely measure the accuracy of multiple tools across varying levels of divergence, we computationally evolved the genome of *E. coli* K-12 W3110 at three different mutation rates: 0.00001 (low), 0.0001 (medium), and 0.001 (high) SNPs per site, per branch. An average of ten rearrangements were introduced, per genome. Each dataset comprises 32 simulated genomes, forming a perfect binary tree.

Approximately 65X coverage of Illumina MiSeq reads was simulated and assembled for each genome to create draft assemblies. For input, the whole-genome alignment programs were given the draft assemblies, and the mapping pipelines the raw reads. Supplementary Figure 1 details the computational performance on the simulated datasets. Parsnp was the only method to finish in fewer than 10 minutes on the 32–genome dataset, with the other methods requiring between 30 minutes to 10 hours. Table 1 gives the accuracy of each tool on each dataset. The tools were benchmarked using true-positive and false-positive rates compared to a known truth, which captures the full alignment accuracy. Figure 1 plots the performance of all tools averaged across all mutation rates.

The whole-genome alignment methods performed comparably across all three mutation rates (Figure 1, red squares), with Mauve exhibiting the highest sensitivity (97.42%) and Parsnp the highest precision (99.99%). Mugsy demonstrated slightly higher sensitivity than Parsnp but with lower precision. Mugsy’s lower precision was traced to a single *fumA*paralog [93] misalignment that generated a high number of false-positive SNPs. All genome alignment methods were affected by misalignment of repeats and missing or low-quality bases in the assembly.

**Table 1.**
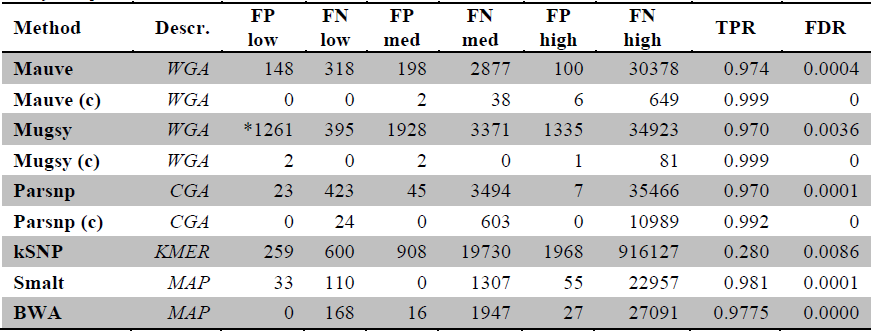
**Core-genome SNP accuracy for simulated *E. coli* datasets**. Data shown indicates performance metrics of the evaluated methods on the three simulated *E. coli* datasets (low, medium and high). Method: Tool used. (c) indicates aligner ran on closed genomes rather than draft assemblies. Descr: Paradigm employed by each method, *WGA* indicates whole-genome alignment, *CGA* indicates core genome alignment, *KMER* indicates k-mer based SNP calls, and *MAP* indicates read mapping. False positive (FP) and false negative (FN) counts for the three mutation rates (low, med, and high). TP: number of SNP calls that agreed with the truth. FP: number of SNP calls that are not in truth set. FN: number of truth SNP calls not detected. True positive rate TPR: TP/(TP+FN). False discovery rate FDR: FP/(TP+FP). A total of 1,299,178 SNPs were introduced into the 32-genome dataset, across all three mutational rates. *Mugsy’s lower precision was traced to a paralog misalignment that resulted in many false-positive SNPs.

**Figure 1.**
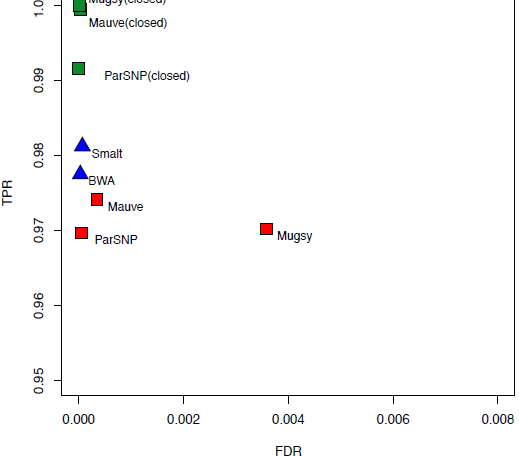
Core-genome SNP accuracy for simulated *E. coli* datasets. Results are averaged across low, medium, and high mutation rates. Red squares denote alignment-based SNP calls on draft assemblies, green squares alignment-based SNP calls on closed genomes, and blue triangles for read mapping. Full results for each dataset are given in Table 1.

**Figure 2.**
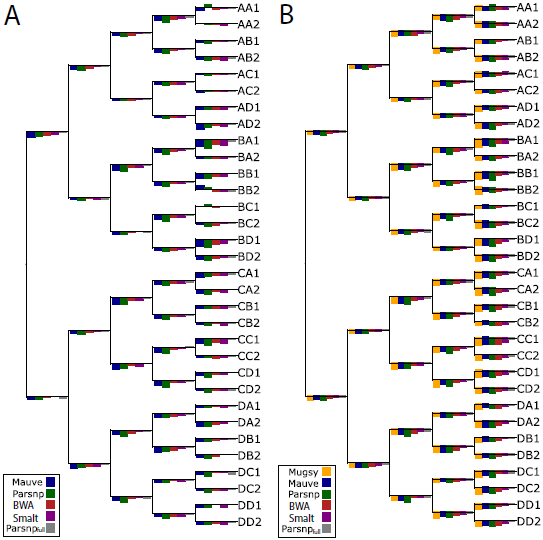
Branch errors for simulated *E. coli* datasets. Simulated *E. coli* trees are shown for medium mutation rate (0.0001 per base per branch). Panel A shows branch length errors as bars, with overestimates of branch length above each branch and underestimates below each branch. Maximum overestimate of branch length was 2.15% (bars above each branch) and maximum underestimate was 4.73% (bars below each branch). Panel B shows branch SNP errors as bars, with false-positive errors above each branch and false-negative errors below each branch. The maximum FP SNP value is 6 (bars above each branch) and maximum FN SNP value is 23 (bars below each branch). Note that the bar heights have been normalized by the maximum value for each tree and are not comparable across trees. Outlier results from Mugsy were excluded from the branch length plot, and kSNP results are not shown. All genome alignment methods performed similarly on closed genomes, with Mauve and Mugsy exhibiting the best sensitivity (Table 1).

Performance of the individual methods was also measured in terms of branch SNP and length error with respect to the true phylogeny (Figure 2). These errors closely followed the false-negative and false-positive rates of each method, with no distinguishable pattern or branch biases. On draft genomes, precise methods such as Parsnp yielded underestimates of branch lengths while more aggressive methods like Mugsy resulted in more overestimates (outliers not shown). The aggressive methods also showed more variance in performance across branches.

#### Comparison on Closed Genomes

Mugsy, Mauve, and Parsnp all performed similarly on finished genomes (Figures 1 and 2, green squares), offering a significant boost in sensitivity over both draft assemblies and reference mapping. Mugsy, Mauve, and Parsnp all exhibited near perfect false-discovery rates (FDR), with Parsnp being the only method to not report a single false positive across the three datasets. Both Mauve and Mugsy were similarly near-perfect in terms of true-positive rates (TPR). The drop in sensitivity (0.9%) for Parsnp on full genomes can be explained by a lack of an LCB extension method. Mugsy was the most affected by draft genomes, going from best on closed genomes to demonstrating more false positives (Table 1) and LCB counts (Table 2) on draft genomes. Parsnp offered the overall best FDR of the genome alignment methods, and the fewest number of LCBs, averaged across both draft and closed genome datasets.

**Table 2.** Comparison of locally collinear alignment block (LCB) count for simulated *E. coli* datasets, on assembled and finished genomes. Method: Tool used. (c) indicates aligner ran on closed genomes rather than draft assemblies. Low: >99.99% similarity, Medium: >99.9% similarity, High: >99% similarity.

| Method | Low | Medium | High |
| --- | --- | --- | --- |
| **Mauve** | 325 | 363 | 519 |
| **Mauve (c)** | 150 | 174 | 333 |
| **Mugsy** | 10977 | 11194 | 16632 |
| **Mugsy (c)** | 237 | 247 | 351 |
| **Parsnp** | 205 | 271 | 344 |
| **Parsnp (c)** | 139 | 190 | 506 |

#### Comparison to Read Mapping Methods

On average, mapping-based methods were as precise and 0.5−1% more sensitive than alignment of draft genomes Figure 1, blue triangles). Smalt showed the highest sensitivity, while BWA was the most specific. The precision of the mapping approaches may be overestimated for this dataset due to the absence of non-core sequence that is known to confound mapping [94]. Parsnp was the only genome alignment method to match the precision of mapping, but with a slight reduction in sensitivity. However, when provided with finished genomes, the whole-genome alignment methods excel in both sensitivity and specificity compared to read mapping. Thus, the performance divide between whole-genome alignment and mapping is entirely due to assembly quality and completeness. Using short reads, both the mapping and assembly-based approaches suffer false-negatives due to ambiguous mappings or collapsed repeats, respectively. Exceeding 99% sensitivity for this test set requires either longer reads (for mapping) or complete genomes (for alignment) to accurately identify SNPs in the repetitive regions.

### Comparison on 31 Streptococcus Pneumoniae Genomess

Parsnp was compared to whole-genome alignment methods using the 31-genome *S.pneumoniae* dataset presented in the original Mugsy publication [36]. Angiuoli and Salzberg compared Mugsy, Mauve, and Nucmer+TBA to measure the number of LCBs and size of the core genome aligned. On this dataset, Parsnp aligns 90% of the bases aligned by Mugsy, while using 50% fewer LCBs (Table 3). In addition, Parsnp ran hundreds of times faster than the other methods, finishing this 31-way alignment in less than 60 seconds.

**Table 3.** Comparison to the 31 *S. pneumoniae* Mugsy benchmark. Time: Method runtime from input to output. Core: Size of the aligned core genome measured in base pairs. LCBs: Number of locally colinear blocks in the alignment.

| Method | Time | Core (bp) | LCBs |
| --- | --- | --- | --- |
| Parsnp | 0.3 min | 1428407 | 1171 |
| Mugsy | 100 min | 1590820 | 2394 |
| Mauve | 377 min | 1568715 | 1366 |
| NUCmer+TBA | 80 min | 1457575 | 27075 |

### Peptoclostridium Difficile Outbreak in the UK

Parsnp and Gingr are particularly suited for outbreak analyses of infectious diseases. To demonstrate this, we applied Parsnp to a recent *P. difficile* outbreak dataset [95]. To generate input suitable for Parsnp, we assembled all genomes using iMetAMOS [96]. It is important to note that this was a resequencing project not intended for assembly and represents a worst case for a core-genome alignment approach; reads ranged from 50 to 100bp in length and some genomes were sequenced without paired ends. The 826-way core genome alignment resulted in 1.4 Gbp being aligned in less than 5 hours. The core genome represented 40% of the *P. difficile* 630 reference genome, consistent with previous findings [97]. Specifically, previous microarray experiments have indicated that 39% of the total CDS in the evaluated *P. difficile* clade pertains to the core genome (1% less than identified by Parsnp). Figure 3 shows a Gingr visualization of the 826-way alignment and clade phylogeny. Related outbreak clusters are immediately visible from the phyletic patterns of the alignment, confirming the primary clades of the tree. In addition, the SNP heatmap highlights the phyletic signature of several subclades, in this case within the known hpdBCA operon [98] that is extremely well conserved across all 826 genomes.

**Figure 3.**
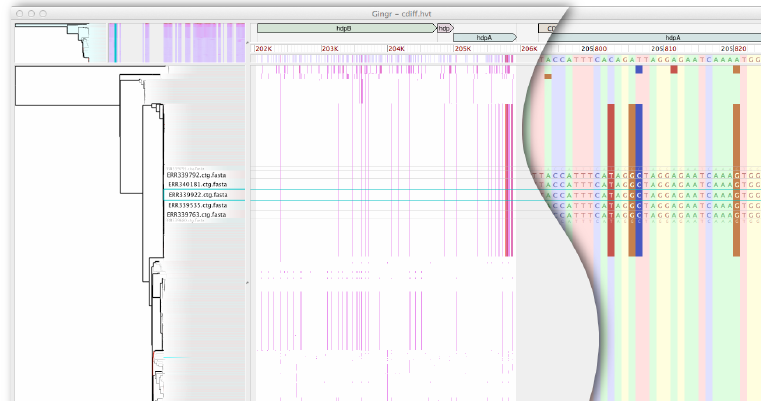
Gingr visualization of 826 *P. difficile* genomes aligned with Parsnp. The leaves of the reconstructed phylogenetic tree (left) are paired with their corresponding rows in the multi-alignment. A genome has been selected (rectangular aqua highlight), resulting in a fisheye zoom of several leaves and their rows. A SNP density plot (center) reveals the phylogenetic signature of several clades, in this case within the fully-aligned hpd operon (hpdB, hpdC, hpdA). The light gray regions flanking the operon indicate unaligned sequence. When fully zoomed (right), individual bases and SNPs can be inspected.

**Figure 4.**
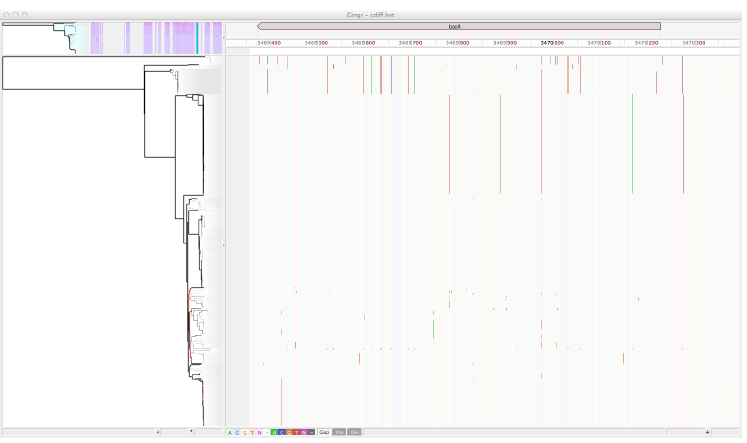
**Conserved presence of *bacA* antiobiotic resistance gene in *P. difficile* outbreak**. Gingr visualization of conserved bacitracin resistance gene within the Parsnp alignment of 826 *P. difficile* genomes. Vertical lines indicate SNPs, providing visual support of subclades within this outbreak dataset.

Figure 4 shows a zoomed view of the 826 *P. difficile* genome alignment in Gingr, highlighting a single annotated gene. Although no metadata is publically available for this outbreak dataset, we identified that *bacA*, a gene conferring antibiotic resistance to bacitracin, is conserved in all 826 isolates. While alternative antibiotic treatments for *P. difficile* infections have been well-studied over the past 20-30 years [99], a recent study reported that 100% of 276 clinical isolates had high-level resistance to bacitracin [100]. In concordance with this study, our results indicate there may be widespread bacitracin resistance across this outbreak dataset. Thus alternative antibiotics, such as vancomycin, could represent better treatment options.

### Mycobacterium Tuberculosis Geographic Spread

For a second case evaluation, we ran Parsnp on a *M. tuberculosis* global diversity dataset [101]. In this case, the raw SNP calls were kindly made available (Iñaki Comas, personal communication), facilitating a direct comparison to the published results. The variant pipeline of Comas *et al.* is similar to our BWA pipeline, but with all SNP calls intersected with MAQ SNPfilter, which discards any SNP with neighboring Indels ±3bp or surrounded by >3 SNPs within a 10bp window. To replicate this study using whole-genome alignment, we assembled all genomes from the raw reads using iMetAMOS and ran Parsnp on the resulting draft assemblies. Figure 5 summarizes the results of the comparison and Figure 6 shows a Gingr visualization of the resulting tree and alignment, with major clades confirmed by correlations in the SNP density display.

**Figure 5.**
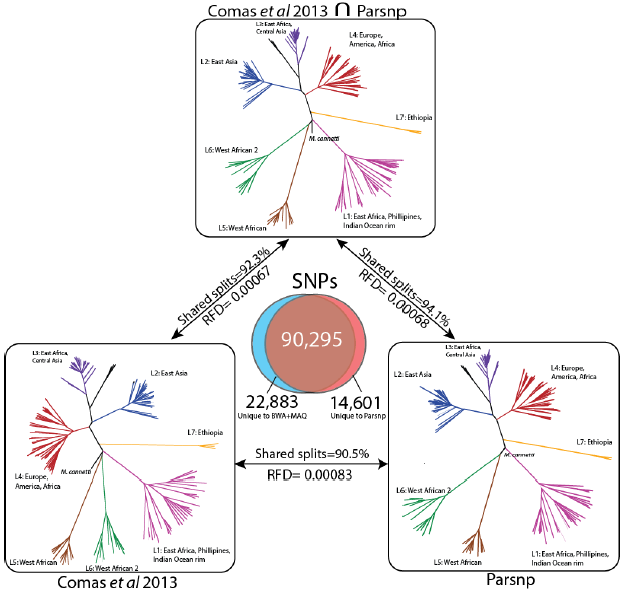
Comparison of Parsnp and Comas *et al.* result on *M. tuberculosis* dataset. A Venn diagram displays SNPs unique to Comas *et al.* [101] (left, blue), unique to Parsnp (right, red), and shared between the two analyses (middle, brown). On top, an unrooted reference phylogeny is given based on the intersection of shared SNPs produced by both methods (90,295 SNPs). On bottom, the phylogenies of Comas *et al.* (left) and Parsnp (right) are given. Pairs of trees are annotated with their Robinson-Foulds distance (RFD) and percentage of shared splits. The Comas *et al.* and Parsnp trees are largely concordant with each other and the reference phylogeny. All major clades are shared and well supported by all three trees.

Given a lack of truth for this dataset, we constructed a reference phylogeny based on the intersection of the Parsnp and Comas *et al.* SNP sets, which excludes potential false positives produced by only one of the methods. We evaluated the accuracy of phylogenetic reconstruction by measuring the Robinson-Foulds distance [102] and calculating the number of shared splits between the resulting trees (Figure 5). The Parsnp generated phylogeny has a higher percentage of shared splits with the reference phylogeny (94.1% versus 92.3% for Comas), while both methods exhibited a similar Robinson-Foulds distance to the reference phylogeny (0.0007).

**Figure 6.**
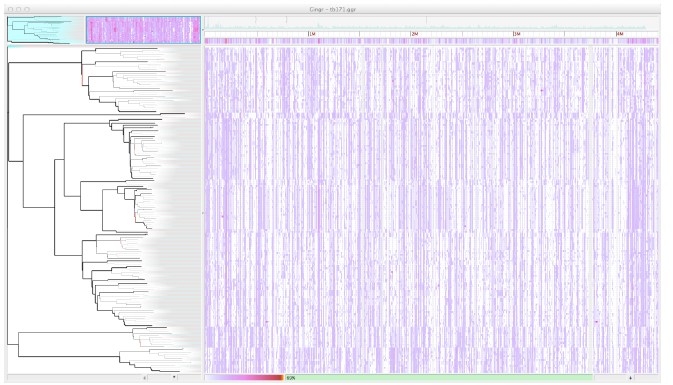
Gingr visualization of 171 *M. tuberculosis* genomes aligned with Parsnp. The visual layout is the same as Figure 3, but unlike Figure 3, a SNP density plot across the entire genome is displayed. Major clades are visible as correlated SNP densities across the length of the genome.

When compared directly, Parsnp was found to share 90,295 of its 104,896 SNPs calls (86%) and 19,838 of its 22,131 SNP positions (90%) with the Comas *et al.* study, resulting in 90.5% shared splits between the reconstructed phylogenies. We further investigated the SNP calls unique to either Parsnp or Comas and found that the majority formed a few well-defined splits that were largely consistent with the reference phylogeny (Supplementary Table 1). These SNPs are likely true positives that were missed by one of the two methods. A smaller fraction of unique SNPs formed single-genome splits, representing potential false positives due to sequencing or mapping error (1,503 for Parsnp, 3,016 for Comas).

### Runtime and Storage Analysis

To evaluate Parsnp’s scalability, we profiled performance across six datasets ranging from 32 genomes to 10,000 genomes. Runtime was observed to increase linearly with additional genomes (Supplementary Figure 2), requiring a few minutes for the 32 genome *E. coli* dataset, 1.5 hours for the 826 genome *P. difficile* dataset, and a maximum of ∼14 hours to align the 10,000 genome set on a 2.2 GHz, 32-core, 1TB RAM server (Supplementary Table 2). In addition, for the 32-genome simulated *E. coli* datasets, Parsnp was 10-100 times faster than all other methods evaluated. Maximum memory usage was 2GB for the 145Mbp *E. coli* dataset and 309GB for the 21Gbp *S. pneumoniae* dataset (Supplementary Table 2). Memory usage can be explicitly limited via a command-line parameter (--max-partition-size) but this results in increased runtime.

In addition to runtime efficiency, Parsnp requires much less storage than the other approaches due to its binary alignment format and the compressive effect of assembly. For the 32-genome *E. coli* dataset, Parsnp’s output totals just 4.5MB, compared to 13GB required to store compressed FASTQ [103] and VCF [104] files and 149MB to store XMFA [38]. Storage reductions are amplified for larger datasets. For example, the raw read data for the *P. difficile* dataset requires 1.4TB of storage (0.6TB compressed). Assembling these data reduces the total to 3.3GB by removing the redundancy of the reads. The XMFA alignment of these assemblies is 1.4GB, and reference-compressed binary format occupies just 15MB. This equates to roughly a 100,000X (lossy) compression factor from raw reads to compressed archive, requiring only 0.08 bits per base to store the full core-genome alignment plus other related information, which is competitive with related techniques like CRAM [105]. As outbreak studies continue to expand in scale, whole-genome assembly and alignment presents a sustainable alternative to the current mapping-based strategies.

## Discussion

Parsnp is orders of magnitude faster than current methods for whole-genome alignment and SNP typing, but it is not without limitations. Parsnp represents a compromise between whole-genome alignment and read mapping. Compared to whole-genome aligners, Parsnp is less flexible because it is designed to conservatively align the core genome and is less sensitive as a result. Compared to read mapping, Parsnp is less robust and requires high-quality assemblies to maximize sensitivity. Thus, the right tool depends on the data and task at hand.

Core-genome alignment and phylogeny reconstruction are critical to microbial forensics and modern epidemiology. When finished or high-quality genomes are available, Parsnp is both efficient and accurate for these tasks. In addition, even for fragmented draft assemblies, Parsnp exhibits a favorable compromise between sensitivity and specificity. Surprisingly, Parsnp matched the specificity of the mapping-based approaches on the simulated datasets. However, multiplexed short-read sequencing followed by mapping still remains the most economical approach for sensitive analysis of large strain collections. Thus, Parsnp is recommended for analyzing high-quality assemblies or when raw read data is not available.

Assembled genomes have a number of advantages over read data—primarily compression and convenience. Storing, sharing, and analyzing raw read datasets incurs significant overhead from the redundancy in sequencing (often 100 fold), and this burden nearly resulted in the closure of the NCBI SRA database [106]. Adding additional orders of magnitude to the already exponential growth of sequencing data is not sustainable. Instead, information in the reads not currently stored in common assembly formats (e.g. allelic variants) should be propagated to the assembled representation, forming a compressed, but nearly lossless format. In this way, genomes could be shared in their native, assembled format, saving both space and time of analysis. Here, we have taken a small step in that direction by identifying low quality bases, as computed by FreeBayes [55]. This allows filtering of low quality and mixed alleles and improves the specificity of the assembly-based approaches. However, more comprehensive, graph-based formats are needed to capture the full population information contained in the raw reads.

Parsnp was also built around the observation that high-quality, finished genome sequences have become more common as sequencing technology and assembly algorithms continue to improve. New technologies, such as PacBio SMRT sequencing [107] are enabling the generation of reference-grade sequences at extremely reduced costs. This presents another opportunity for Parsnp—the construction and maintenance of core genomes and trees for clinically important species. With well defined reference cores, outbreaks could be accurately typed in real-time by mapping sequences directly to the tree using phylogenetically aware methods such as pplacer [108] or PAGAN [109]. Such a phylogenetic approach would be preferable to alternative typing schemes based on loosely defined notions of similarity, such as pulse-field electrophoresis (PFGE) [110] and multi-locus sequence typing (MLST) [111].

## Conclusion

Parsnp offers a highly efficient method for aligning the core genome of thousands of closely related species, and Gingr provides a flexible, interactive visualization tool for the exploration of huge trees and alignments. Together, they enable analyses not previously possible with whole-genome aligners. We have demonstrated that Parsnp provides highly specific variant calls, even for highly fragmented draft genomes, and can efficiently reconstruct recent outbreak analyses including hundreds of whole genomes. Future improvements in genome assembly quality and formats will enable comprehensive cataloging of microbial population variation, including both point and structural mutations, using genome alignment methods such as Parsnp.

## Materials and Methods

### Software and Configurations

Mugsy [36] v1.23 and Mauve Aligner [31,33] v2.3.1 were run using default parameters on assembled sequences. mauveAligner was selected instead of progressiveMauve due to improved performance on the simulated *E. coli* datasets, which do not contain subset relationships. kSNP v2.0 [67] was run with a k-mer size of 25 on both the raw read data and the assemblies; the assemblies were merged with Ns using the provided `merge_fasta_contigs.pl` utility. Raw MAF/XMFA/VCF output was parsed to recover SNPs and build MultiFASTA files using a conversion script [112].

Smalt version 0.7.5 was run with default parameters for paired reads, mirroring the pipeline used in several recent SNP typing studies [92,113–115]. `Samtools view` was used to filter for alignments with mapping qualities greater than or equal to 30. Variants were called by piping `samtools mpileup` output into `bcftools view` with the -v (variants only), -g (genotype) and -I (skip Indels) flags. Variants were then filtered with `VCFUtils varFilter` with the -d (minimum read depth) parameter set to 3. Variants for all samples of each set were called concomitantly by providing `samtools mpileup` with all BAM files.

BWA [53] was run in its standard paired-end alignment mode with default parameters, using `aln` to align each set of ends and `sampe` to produce a combined SAM file. `Samtools view` was used to filter for alignments with mapping qualities greater than or equal to 30. Variants were called by piping `samtools mpileup` output into `bcftools` view with the -v (variants only), -g (genotype) and -I (skip Indels) flags. Variants were then filtered with `VCFUtils varFilter` with the -d (minimum read depth) parameter set to 3. As with Smalt, variants for all samples of each set were called concomitantly by providing `samtools mpileup` with all BAM files.

FastTree v2 [90] was used to reconstruct phylogenies using default parameters.

### E. *Coli* K-12 W3110 Simulated Dataset

The complete genome of *E. coli* K-12 W3110 [116], was downloaded from RefSeq (AC_000091). This genome was used as the ancestral genome and evolution was simulated along a balanced tree for three evolutionary rates using the Seq-Gen package [117] with parameters `mHKY -t4.0 -l4646332 -n1 -k1` and providing the corresponding binary tree evolved at three evolutionary rates: 0.00001, 0.0001, and 0.001 SNPs per site, per branch. This corresponds to a minimum percent identity of approximately 99%, 99.9%, and 99.99% between the two most divergent genomes, respectively, reflecting the variation seen in typical outbreak analyses. No small (< 5bp) or large Indels were introduced, but an average of ten 1Kbp rearrangements (inversions and translocations) were added, per genome, using a custom script. Paired reads were simulated to model current MiSeq lengths (2x150bp) and error rates (1%). Moderate coverage, two million PE reads (64X coverage), was simulated for each of the 32 samples using wgsim (default parameters, no Indels), from samtools package version 0.1.17 [56].

Two of the simulated read sets were independently run through iMetAMOS [96] to automatically determine the best assembler. The consensus pick across both datasets was SPAdes version 3.0 [83], which was subsequently run on the remaining 30 simulated read sets using default parameters. The final contigs and scaffolds files were used as input to the genome alignment methods. For mapping methods, the raw simulated reads were used. For accuracy comparisons, Indels were ignored and called SNPs were required to be unambiguously aligned across all 32 genomes (i.e. not part of a subset relationship; SNPs present but part of a subset relationship were ignored).

### S. Pneumoniae Dataset

A full listing of accession numbers for the 31-genome *S. pneumoniae* dataset is described in [36]. For scalability testing, *Streptococcus pneumoniae* TIGR4 (NC_003028.3) was used to create a pseudo-outbreak clade involving 10,000 genomes evolved along a star phylogeny with on average 10 SNPs per genome.

### M. Tuberculosis Dataset

We downloaded and assembled sequencing data from a recently published study of *M. tuberculosis* [101]. 225 runs corresponding to project ERP001731 were downloaded from NCBI SRA and assembled using the iMetAMOS ensemble of SPAdes, MaSuRCA, and Velvet. The iMetAMOS assembly for each sample can be replicated with the following commands, which will automatically download the data for `RUN_ID` directly from SRA:

> `initPipeline -d asmTB -W iMetAMOS -m RUN_ID -i 200:800`
> `runPipeline -d asmTB -a spades,masurca,velvet -p 16`

The *M. tuberculosis* dataset included a mix of single and paired-end runs with a sequence length ranging from 51-108bp. The average k-mer size selected for unpaired data was 26, resulting in an average of 660 contigs and an N50 size of 17Kbp. For paired-end data, the average selected k-mer was 35, resulting in an average of 333 contigs and an N50 size of 43Kbp. Assemblies containing more than 2,000 contigs, or 1.5X larger/smaller than the reference genome, were removed. The final dataset was reduced to 171 genomes, limited to labeled strains that could be confidently matched to the strains used in the Comas *et al.* study for SNP and phylogenetic comparison.

### P. Difficile Dataset

Note, *Clostridium difficile* was recently renamed to *Peptoclostridium difficile [118]*. We downloaded and assembled sequencing data from a recently published study of *P. difficile* [95]. 825 runs corresponding to project ERP003850 were downloaded from NCBI SRA [88] and assembled within iMetAMOS this time only using SPAdes, which was identified as the best performer on the *M. tuberculosis* dataset. The iMetAMOS assembly for each sample can be replicated with the following commands, which will download the data for `RUN_ID` directly from SRA:

> `initPipeline -d asmPD -W iMetAMOS -m RUN_ID -i 200:800`
> `runPipeline -d asmPD -a spades -p 16`

The *P. difficile* dataset included paired-end runs with a sequence length ranging from 51-100bp. SPAdes was selected as the assembler and run with k-mer sizes of 21, 33, 55, and 77. The assemblies had an average of 660 contigs and an N50 size of 138Kbp. Assemblies containing more than 2,000 contigs, or 1.5X larger/smaller than the reference genome, were removed.

## Abbreviations

Bp: base pair; Mbp: million base pairs; LCB: locally collinear block; MUM: maximal unique match; MUMi: similarity index based on maximal unique matches; SNP: single-nucleotide polymorphism; Indel: insertion or deletion; ERA: European Read Archive; SRA: Sequence Read Archive; PE: paired-end; NGS: Next generation sequencing; VCF: variant call format; XMFA: extendend multi-fasta format.

## Data and Software Availability

All data, supplementary files, assemblies, packaged software binaries and scripts described in the manuscript are available from [112]. Source code of the described software, including Parsnp and Gingr, is available for download from: http://github.com/marbl/harvest

## Authors’ Contributions

TJT, BDO, and AMP conceived the method, designed the experiments, and drafted the manuscript. TJT implemented Parsnp and BDO implemented Gingr. TJT, BDO, and SK performed the experiments. All authors read and approved the final manuscript.

## Acknowledgments

We would like to thank Iñaki Comas for providing the raw data for the TB study; Xiangyu Deng for helpful comments on the manuscript and software; and Bradd Haley, Bill Klimke, Jason Sahl, Maliha Aziz, and Suzanne Bialek-Davenet for feedback on early versions of the software. This work was funded under Agreement No. HSHQDC-07-C-00020 awarded by the Department of Homeland Security Science and Technology Directorate (DHS/S&T) for the management and operation of the National Biodefense Analysis and Countermeasures Center (NBACC), a Federally Funded Research and Development Center. The views and conclusions contained in this document are those of the authors and should not be interpreted as necessarily representing the official policies, either expressed or implied, of the U.S. Department of Homeland Security. In no event shall the DHS, NBACC, or Battelle National Biodefense Institute (BNBI) have any responsibility or liability for any use, misuse, inability to use, or reliance upon the information contained herein. The Department of Homeland Security does not endorse any products or commercial services mentioned in this publication.

