## Supplementary Material for "Rapid Core-Genome Alignment and Visualization for Thousands of Intraspecific Microbial Genomes"

### Supplementary Materials

- **Supplementary Table 1**. SNPs unique to each method characterized by the five most common splits found.
- **Supplementary Table 2**. Performance profile of Parsnp runtime (MUM+alignment) on all evaluated datasets.
- **Supplementary Figure 1**. Runtime comparison for the whole-genome alignment methods on the simulated 32-genome *E. coli* W3110 dataset.
- **Supplementary Figure 2**. Timing performance from 32 to 10,000 *S. pneumoniae* genomes.

**Supplementary Table 1**. SNPs unique to each method characterized by the five most common splits represented. Combined, these five splits account for approximately half of the unique SNPs and ~70% of the unique SNP positions. **Col Cnt:** the number of alignment columns (i.e. SNP positions) supporting the split**. Split:** The identified split overlaid on the reference tree, with the highlighted genomes forming one half of the split. **SNP count:** the total number of SNPs that pertain to the split.

| Method | Col Cnt | Split | SNP count |
|---|---|---|---|
| **Parsnp** | 28 (1%) | 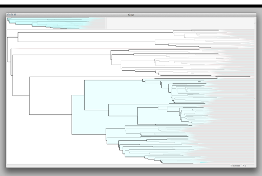 | 3052 (21%) |
| **Parsnp** | 129 (6%) | 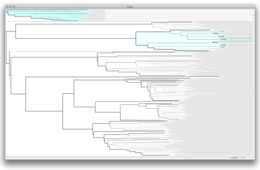 | 1677 (11%) |
| **Parsnp** | 1503 (66%) | Single genome splits | 1503 (10%) |
| **Parsnp** | 23 (1%) | 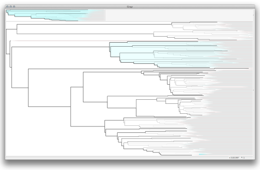 | 1127 (8%) |
| **Parsnp** | 8 (<1%) | 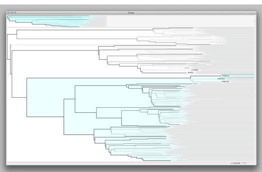 | 896 (6%) |

| | | | |
|---|---|---|---|
| **Comas** | 30 (1%) | 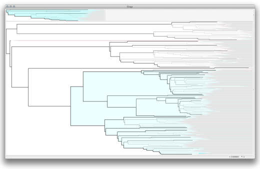 | 3270 (14%) |
| **Comas** | 3016 (63%) | Single genome splits | 3016 (13%) |
| **Comas** | 74 (2%) | 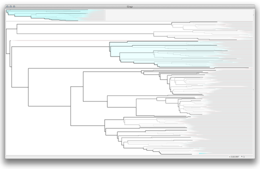 | 2516 (11%) |
| **Comas** | 109 (2%) | 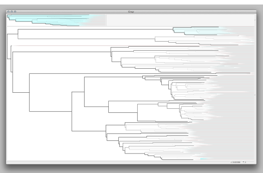 | 1199 (5%) |
| **Comas *et al*** | 32 (1%) | 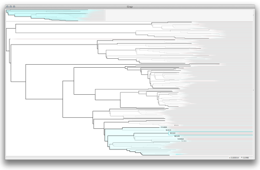 | 1127 (5%) |

**Supplementary Table 2**. Performance profile of Parsnp runtime (MUM+alignment) on all evaluated datasets. Results were generated on a 32-core, 2.2GHz, 1TB RAM Linux server. **Dataset:** the genome set. **Num Genomes:** the number of genomes aligned. **Aligned:** total Mbp aligned. **MUM:** the time spent finding maximal unique matches. **MUSCLE:** the time spent performing gapped multi-alignment with MUSCLE. **Total:** total Parsnp runtime (sum of MUM and MUSCLE). **Mem:** maximum memory usage.

| Dataset | Num Genomes | Aligned (Mbp) | MUM (min) | MUSCLE (min) | Total (min) | Mem (GB) |
|---|---|---|---|---|---|---|
| ***E. coli* (avg)** | 32 | 142 | 2 | 2 | 4 | 2 |
| ***M. tuberculosis*** | 171 | 424 | 12 | 20 | 32 | 14 |
| ***C. dificile*** | 826 | 1,392 | 46 | 39 | 85 | 71 |
| ***S. aureus SIM*** | 10,000 | 21,000 | 668 | 201 | 869 | 309 |

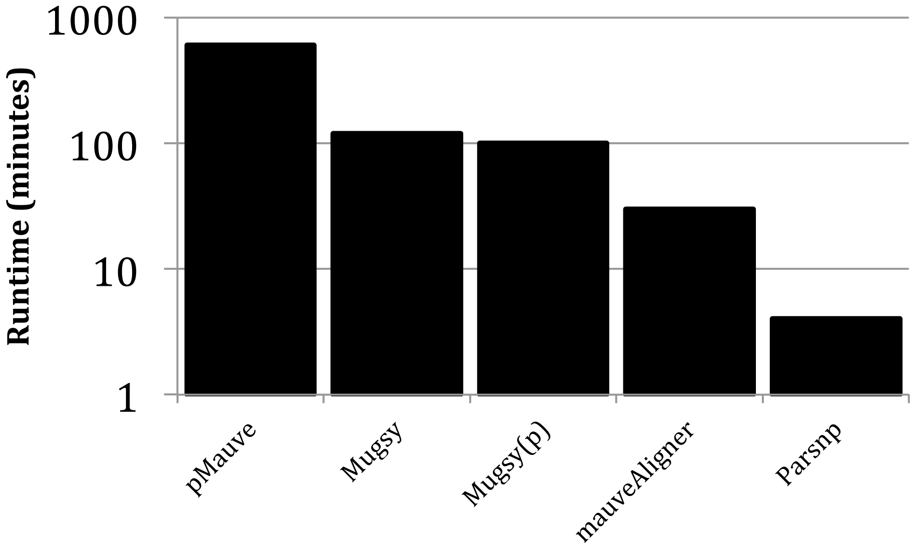


**Supplementary Figure 1**. Runtime comparison for the whole-genome alignment methods on the simulated 32-genome *E. coli* W3110 dataset. The y-axis is log scale. pMauve = progressiveMauve, and Mugsy(p) indicates Mugsy with a parallelized NUCmer search. All programs were allocated 32 cores on the hardware noted above. Note that Mugsy is not multithreaded.

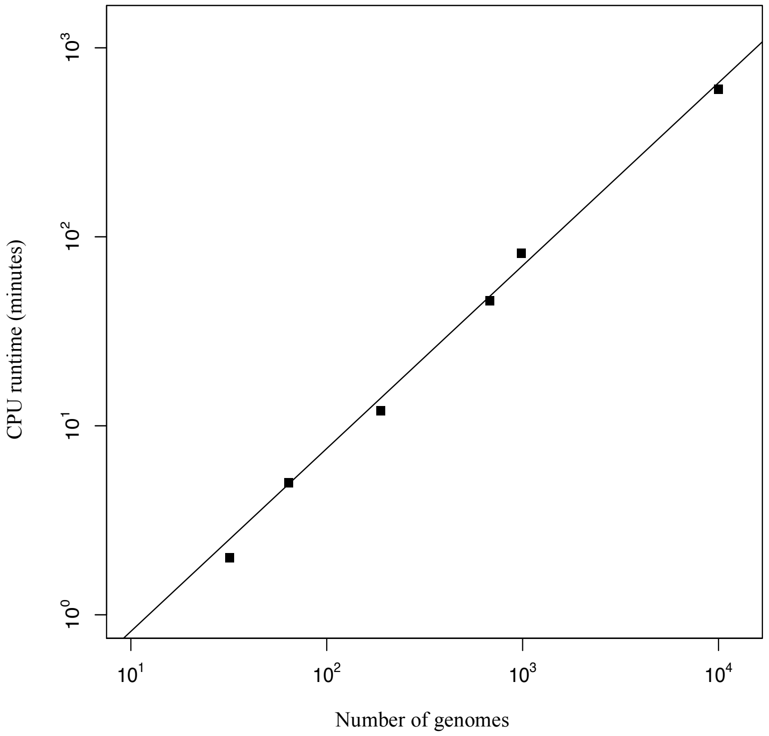


**Supplementary Figure 2.** Timing performance from 32 to 10,000 *S. pneumoniae* genomes. The x-axis indicates the number of genomes and the y-axis the wall clock time for core-genome alignment. Alignments were performed on the hardware noted above. The gray line represents a linear time relationship between the number of genomes and search time.
